# Tumorigenic epithelial clusters are less sensitive to moderate osmotic stresses due to low amounts of junctional E-cadherin

**DOI:** 10.1101/2020.10.26.355891

**Authors:** Danahe Mohammed, Park Young Chan, Jeffrey J. Fredberg, David A. Weitz

## Abstract

The migration of tumorigenic epithelial cells is a critical step for metastatic breast cancer progression. Although the role of the extracellular matrix in breast cancer cell migration has been extensively described, the effect of osmotic stress on the migration of tumor breast epithelial cohorts remains unclear. Most of our understanding on the effect of osmotic stresses on cell migration comes from studies at the level of the single cell in isolation and does not take into account cell-cell interactions. Here, we study the impact of moderate osmotic stress on the migration of epithelial clusters composed of either non-tumorigenic or tumorigenic epithelial cells. We observe a decrease in migration distance and speed for non-tumorigenic epithelial cells but not for tumorigenic ones. To explain these differences, we investigate how osmotic stress impacts the mechanical properties of cell clusters and affects cell volumes. After application of osmotic stress renal epithelial cells become stiffer whereas non-tumorigenic and tumorigenic breast epithelial cells do not. In addition, tumorigenic cells are shown to be less sensitive to osmotic stress than non-tumorigenic cells, and this difference is associated with lower levels of E-cadherin expression. Using EGTA treatments, we confirm that the establishment of cell-cell adhesive interactions is a key component of the behavior of epithelial clusters in response to osmotic stress. This study provides evidence on the low sensitivity of tumorigenic epithelial clusters to moderate osmotic stress and highlights the importance of cadherin-based junctions in response to osmotic stress.

## Introduction

A fundamental challenge is to understand how cell mechanical properties are impacted by the microenvironment. Recent research shows that mechanical properties of cells and the tumor microenvironment can influence cell behavior, tumor growth and dynamics, in ways that remain poorly understood (La Porta et al., 2015; Helmlinger et al., 1997; Cheng et al., 2009; Samuel et al., 2011; Montel et al., 2011; Tse et al., 2012; Dumitru et al.). Cell behavior can be influenced by the matrix rigidity (Hoffman et al., 2011; Riaz et al., 2016; Versaevel et al., 2014), the confinement (Mohammed et al., 2019a; Lantoine et al., 2016; Alaimo et al., 2020; Miermont et al., 2013) or the topography (Mohammed et al., 2019b; Charras and Paluch, 2008; Mohammed et al., 2020). In the case of cancer, the properties of the tumor matrix are also impacted by the environment which determines how cancer cells polarize, adhere, contract and migrate, thereby regulating their invasiveness (Xu et al., 2012). One of the important parameters of the microenvironment for cell regulation is osmotic pressure (Lucké and McCutcheon, 1932; Dick, 1959; Fritz, 1986). The osmotic pressure is determined by the concentration of ions and proteins within the cell microenvironment. Controlling osmotic pressure has various impacts on cell behavior, such as modification of the volume and morphology as well as changes in cell motility (Carton et al., 2003; Karlsson et al., 2013; Lagana et al., 2000; Lewis et al., 2002; Stroka et al., 2014). Moreover, major changes in the functioning of cells are observed (Racz et al., 2007; Nielsen et al., 2008; Miermont et al., 2013; Guo et al., 2017). Hyperosmotic stress alters a variety of cellular processes, for example, it disrupts the cytoskeleton structure (Miermont et al., 2013; Chowdhury et al., 1992; Slaninová et al., 2000), induces chromatin remodeling (Kültz and Chakravarty, 2001; Mas et al., 2009), and triggers cell cycle arrest (Escoté et al., 2004; Clotet et al., 2006) and apoptosis (Silva et al., 2005; Galvez et al., 2001). In a pathological context, recent studies have shown that serum albumin, whose main function is to regulate the osmotic pressure of blood, is associated with better survival during cancer (Miermont et al., 2013; Seaton, 2001). The transmigration capability of cancer cells, which enables cancer cells to invade surrounding tissue, decreases when a low osmotic pressure is applied (~1 kPa) (La Porta et al., 2015). In addition, osmotic pressure has an impact on stem cell fate, suggesting that changes in cell volume due to osmotic pressure alter physiological parameters like stem-cell differentiation (Guo et al., 2017). Taken together, these observations have led to the recognition of osmotic pressure as an important parameter in physiological and pathological situations and demonstrate a clear link among cell behavior, cancer and osmotic pressure. However, most previous work used levels of osmotic stress (Racz et al., 2007; Nielsen et al., 2008; Miermont et al., 2019; Bounedjah et al., 2012; Ignatova and Gierasch, 2006) far from the physiological and pathological ranges (Kodama and Mori, 2019; Miermont et al., 2013; Le et al., 2006; Havard et al., 2011; McGrail et al., 2015). Thus, the mechanism by which osmotic stress modulates epithelial cell migration and the precise impact it has on tumorigenic epithelial cells remains poorly understood. Further, most previous studies of the role of osmotic stress were performed at the single-cell level (La Porta et al., 2015; Guo et al., 2017) and do not take into account the modulation of cell-cell adhesions among tumor epithelial cells. Cell-cell interactions are crucial for tissue integrity, collective cell migration and cancer propagation. In this paper, we determine the effect of moderate osmotic stresses on the migration of minimal cell clusters and investigate the role of cell-cell junctions. To apply hyperosmotic stress, we add 300-Da polyethylene glycol (PEG 300) directly to the medium. We demonstrate that when non-tumorigenic epithelial cells are cultured under osmotic stress, not only the cell volume but also cell motility decrease. By contrast, this behavior is not observed with tumorigenic epithelial cells, which are shown to be less affected by osmotic pressure. By inhibiting cell-cell adhesions in non-tumorigenic cells we show that they become less affected by osmotic stresses, similar to what is observed in tumorigenic cells which have lower cell-cell adhesions. These observations reveal a surprising and previously unidentified role of cell-cell adhesions in the regulation of cell volume and migration in response to variations of osmotic pressure.

## Results

### Osmotic stress modulates migration distance and speed of non-tumorigenic epithelial cells but not tumorigenic breast epithelial cells

To investigate whether osmotic pressure can affect the migration of breast epithelial cells, we cultivate three epithelial cell lines on glass substrates coated with fibronectin (FN). We use a renal epithelial cell line (MDCK, Fig. 1A), a non-tumorigenic epithelial breast line (MCF-10A, Fig. 1D) and a tumorigenic epithelial breast line (MDA-MB-231, Fig. 1G). After 8 hours in culture with fresh media, we add three different concentrations of PEG 300 (Figure S1) into the culture medium to exert well-controlled moderate osmotic stresses on cell cultures ranging from ~128 to ~231 kPa, as observed in physiological situations (Kodama and Mori, 2019; Miermont et al., 2013; Le et al., 2006; Havard et al., 2011; McGrail et al., 2015). The motility of the different epithelial cell lines in response to osmotic stresses is then observed with time-lapse microscopy. For each cell line, experiments are started with an initial acquisition of 8 hours under isotonic conditions (culture medium), representing the control. Then a PEG solution of 1%, 1.5% or 2.5% (v/v) is added to the culture medium and an additional time-lapse acquisition of 8 hours is collected. To avoid any artefacts and to ensure the lowest variability in our analysis, the migration distance and speed determined for each PEG concentration are normalized by the values obtained from the control experiments performed on the same culture. This normalization enables us to determine the variation of migration distance and speed from the control for each PEG concentration. We find that the maximum distance from the origin (Fig. 1B) and the average migration speed (Fig. 1C) of MDCK cells decreases significantly in response to moderate osmotic stress. The normalized distance and average speed decrease monotonically with increasing PEG concentration, reaching values of 0.40 +/− 0.13 μm (n=7) and 0.68 +/−0.07μm/min (n= 7) respectively at a concentration of 2.5% (v/v). The higher the PEG concentration, the lower the migrating distance and the speed. Interestingly, our results indicate that normalized migration distance (Fig. 1E) and speed (Fig. 1F) of MCF-10A cells are significantly decreased by moderate osmotic stresses, independent of the PEG concentration. The migration parameters of MCF-10A are not statistically affected by the PEG concentration, suggesting that MCF-10A are sensitive to very low osmotic stress (130 kPa). Surprisingly, our findings demonstrate that the migration parameters of MDA-MB-231 cells are not affected by osmotic stresses for PEG concentrations ranging from 1% to 2.5% (v/v) (Figs. 1H and1I). Interestingly, the differences in migratory distance (Fig. 1J) and speed (Fig. 1K) are statistically different at 2.5% (v/v) PEG for the three cell lines. These results demonstrate that tumorigenic epithelial breast cells are more resistant to osmotic stress changes than are MDCK and non-tumorigenic epithelial cells.

**Figure 1.**
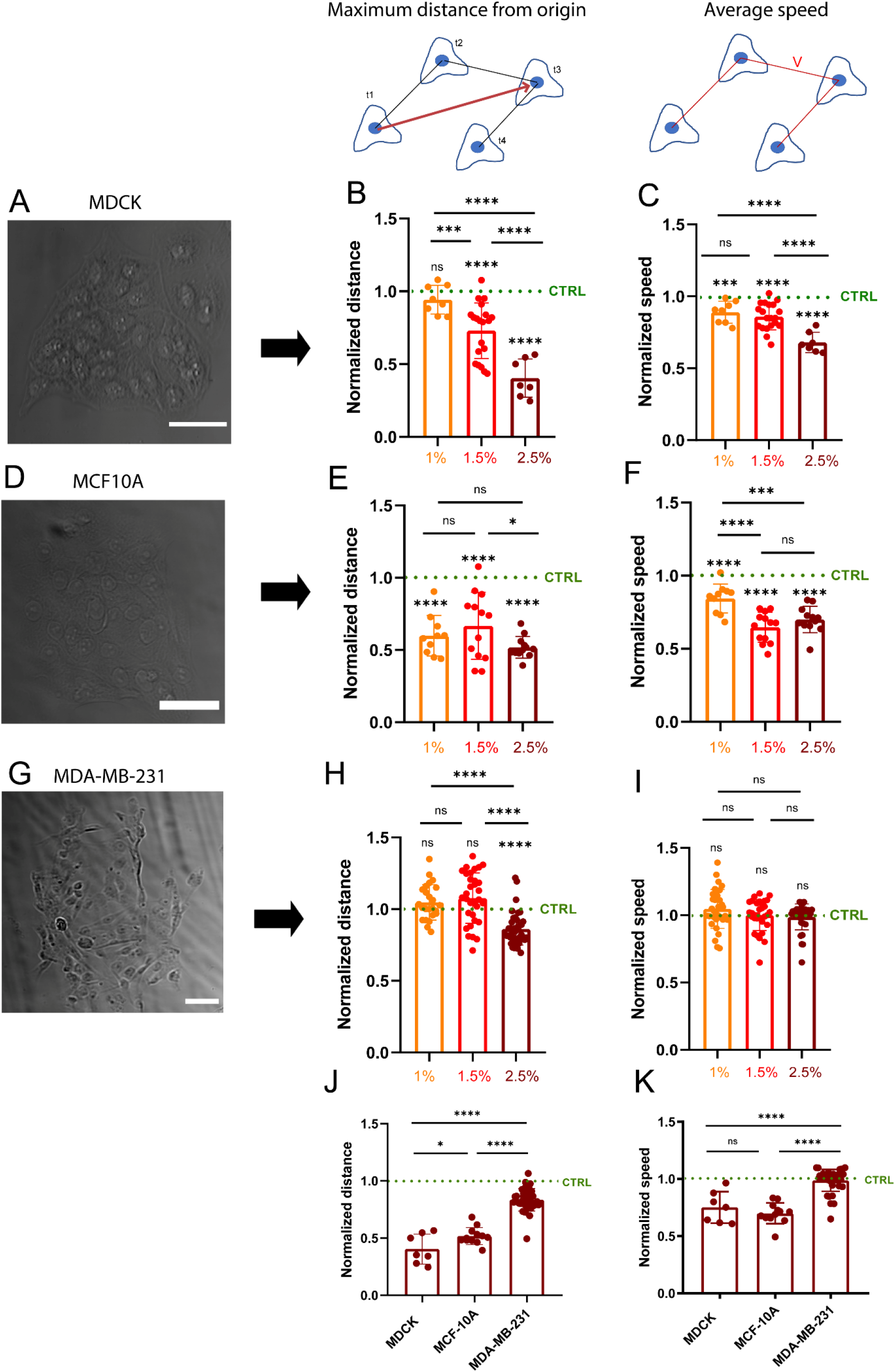
Osmotic-stress modulates migration and speed of non-tumorigenic epithelial cells, but not those of tumorigenic breast epithelial cells. Brightfield images of (A) MDCK, (D) MCF-10A and (G) MDA-MB-231 epithelial cell clusters. Scale bars correspond to 50 μm. Normalized migration distances of (B) MDCK, (E) MCF-10A and (H) MDA-MB-231 treated with 1% (yellow), 1.5% (red) and 2.5% (dark red) PEG concentrations. For each cell type and PEG concentration, maximal distances of migration are normalized by maximum distance of migration of control cells. Normalized migration speeds of (C) MDCK, (F) MCF10A and (I) MDA-MB-231 treated with 1% (yellow), 1.5% (red) and 2.5% (dark red) PEG concentrations. For each cell type and PEG concentration, migration speeds are normalized by migration speeds of control cells. (J) Normalized migration distances and (K) normalized migration speeds for MDCK, MCF-10A and MDA-MB-231 treated with 2.5% PEG. 7≤ n ≤20 for MDCK, 10≤ n ≤13 for MCF10A and 24≤n≤40 for MDA-MB-231. *0.01<p<0.05, **0.001<p<0.01, ***0.0001<p<0.001, ****p<0.0001 and n.s. non-significant.

### The rigidity of tumorigenic breast epithelial cells is not affected by osmotic stresses

To help understand the differences in migration among MDCK, non-tumorigenic (MCF10A) and tumorigenic epithelial breast cells (MDA-MB-231) in response to moderate osmotic stress, we measure the change in cell stiffness of the three cell lines in response to osmotic stress at 1.5% and 2.5% (v/v) using optical magnetic twisting cytometry (OMTC)(Zhou et al., 2009) (Fig. 2A). Similar to the migration parameters, we normalize the cell stiffness in response to 1.5% and 2.5% (v/v) PEG with the baseline stiffness measured on the same cells in the absence of PEG. We observe that MDCK cells (Fig. 2B) are stiffer when subjected to osmotic stress, whereas the normalized cell stiffness of MCF-10A (Fig. 2C) and MDA-MB-231 (Figs. 2D) cells remains constant, independent of osmotic stress. These results suggest that the lower migration efficiency of MDCK is associated with cell stiffening, whereas the tumor epithelial cells maintain a constant cell stiffness and hence a constant migratory behavior (Fig. 2E).

**Figure 2.**
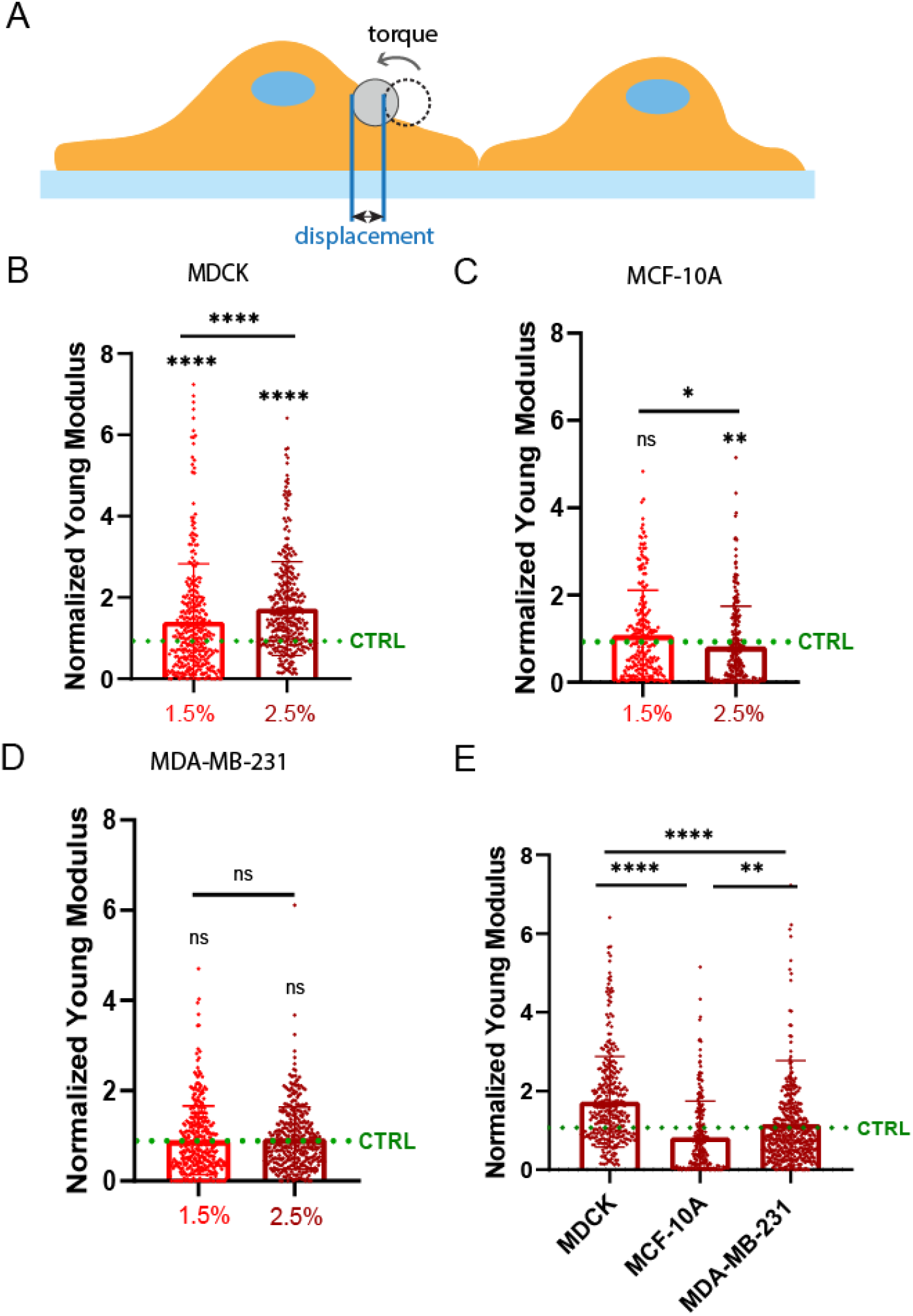
The rigidity of tumorigenic breast epithelial cells is not affected by osmotic stress. (A) Schematic representation of OMTC measurements. Elastic modulus with different concentrations of PEG 1.5% (red) and 2.5% (dark red) normalized by those under isotonic condition for (B) MDCK, (C) MCF-10A and (D) MDA-MB-231 cells. (E) Comparison of normalized elastic modulus for the three cell lines exposed to 2.5% PEG with 70≤n≤140 for all experiments. *0.01<p<0.05, **0.001<p<0.01, ****p<0.0001 and n.s. non-significant.

### Moderate osmotic stress leads to a reduction in cell volume, but tumorigenic breast epithelial cells are less sensitive to osmotic stress

To further investigate how moderate osmotic stresses act on cell stiffness, we use confocal microscopy to measure the volume of minimal tissues grown on circular FN micropatterns and composed of either MDCK, MCF-10A or MDA-MB-231 cells. When single, isolated cells are subjected to an external osmotic pressure, their volume decreases due to water efflux, and their stiffness increases(Guo et al., 2017). The internal osmotic pressure of a cell is regulated by the concentration of ions and small proteins and must adapt to changes in the external osmotic pressure to eliminate any gradient in pressure across the cell membrane. This can result in a significant change in subcellular macromolecular density. To standardize our measurements, we form minimal epithelial tissues of MDCK (Fig. 3A), MCF10A (Fig. 3B) and MDA-MB-231 (Fig. 3C) by using circular micropatterns of FN of 200 μm diameter. The FN micropatterns enable control of the size of the minimal tissues formed and retain their cell density constant, thus limiting the variability between epithelial clusters. By labeling the cytoplasm in live conditions with cell tracker, we determine the cell volume of the whole circular tissues under isotonic conditions using culture medium, and in response to moderate osmotic stresses of 1%, 1.5% and 2.5% (v/v) of PEG. We normalize the volumes of the cells subjected to PEG by those of the control for each cell line to highlight variations of cell volume. Osmotic stress leads to significant decreases in volume for MDCK (~73%, Fig. 3D) and MCF10-A (~46%, Fig. 3E), whereas only a moderate decrease in volume is observed for tumorigenic breast epithelial cells, MDA-MB-231 (~29%, Fig. 3F). Thus, MDA-MB-231 cells are less sensitive to moderate osmotic stress than are MDCK and MCF-10A cells, suggesting that tumorigenic epithelial cells are more robust to variations of osmotic pressure than non-tumorigenic epithelial cells.

**Figure 3.**
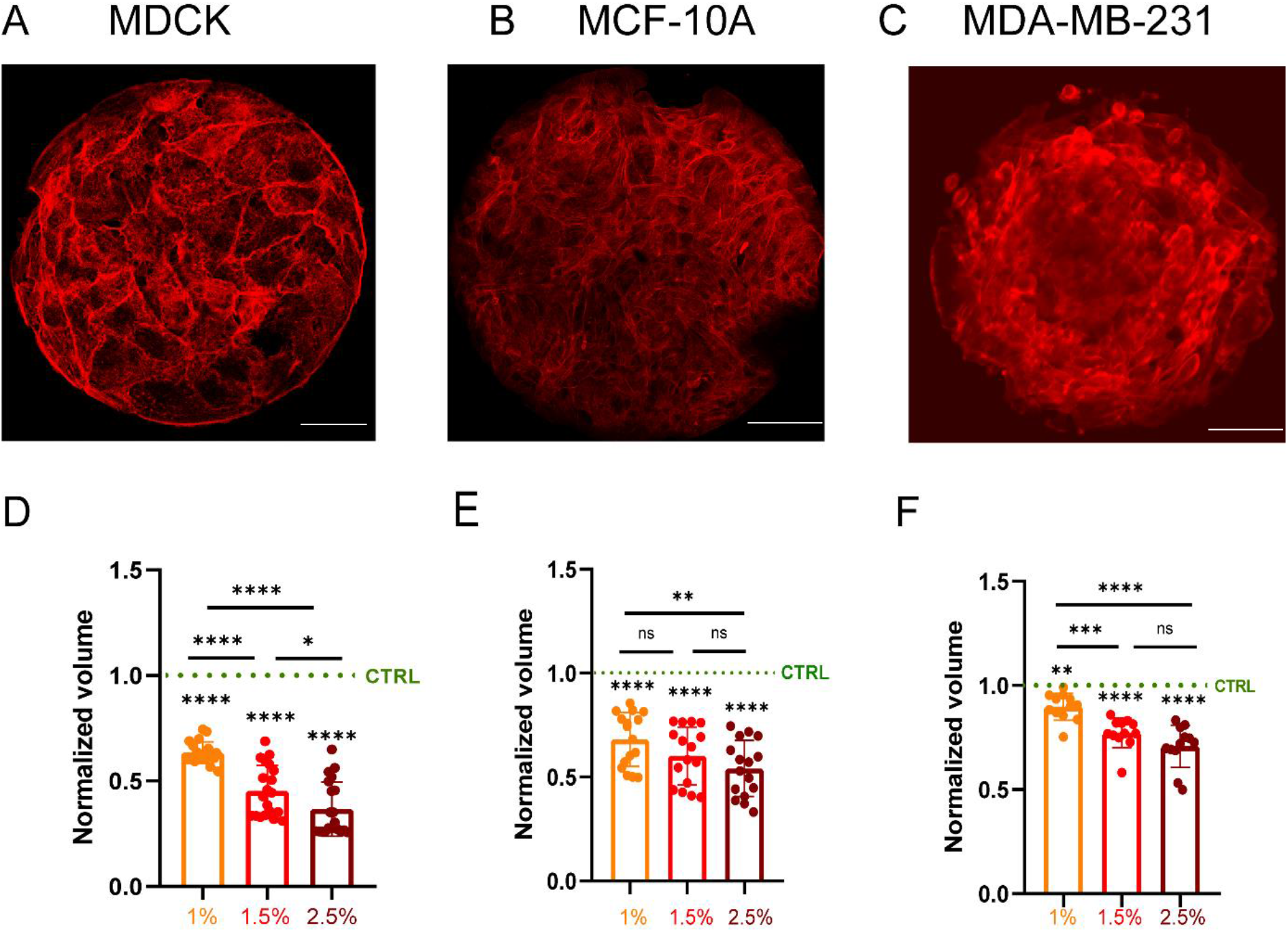
Moderate osmotic stress leads to a loss of cell volume, but tumorigenic epithelial cells are less sensitive than MDCK and non-tumorigenic cells. Immunofluorescence images of (A) MDCK, (B) MCF-10A and (C) MDA-MB-231 minimal tissues grown on circular FN micropatterns with 100 and 200 μm diameter. Actin is stained in red. Scale bar corresponds to 10 μm for MDCK and 40 μm for MCF-10A and MDA-MB-231. Normalized volumes of (D) MDCK, (E) MCF-10A and (F) MDA-MB-231 micropatterned tissues treated with three different PEG concentrations: 1% (yellow), 1.5% (red) and 2.5% (dark red). For each cell type, the volumes of PEG-treated tissues are normalized with the volume of micropatterned tissues. n=20 (MDCK), n=16 (MCF10A), n=12 (MDA-MB-231). *0.01<p<0.05, **0.001<p<0.01, ***0.0001<p<0.001, ****p<0.0001 and ns non-significant.

### The establishment of cell-cell interactions leads to a larger volume loss in response to osmotic stresses

To understand the mechanism that lead to large variations of volume loss in minimal epithelial clusters of MDCK, MCF10A and MDA-MB-231 cells, we investigate the role of cell-cell adhesive interactions. To accomplish this, we treat cells with EDTA which disrupts cell-cell junctions^45-47^ and thus converts the micropatterned cluster of cells into a group of isolated cells. Interestingly, our results show that the PEG induced volume loss of cells treated with EGTA is statistically the same for MDCK (~19%, Fig. 4A), MCF10A (~17%, Fig. 4B) and MDA-MB-231 (~19%, Fig. 4C) cells. This suggests that the establishment of cell-cell adhesive interactions in the clusters of cells may be important in causing the differences in the volume loss under osmotic stresses for the different cell lines. Interestingly, for tumorigenic cell line, there is a smaller difference in cell volume loss between single cells and cell clusters (Fig. 4C). To further understand these results, we determine the contrast in E-cadherin between the junction and the cytoplasm in the three cell types by immunostaining experiments (Fig. 4D). We observe that MDCK tissues exhibit a larger amount of E-cadherins at the cell-cell junctions than do MCF-10A and MDA-MB-231 (Fig. 4E). Interestingly, E-cadherins are more localized in the cytoplasm in the MDA-MB-231 cell clusters. We observe a linear correlation between the volume loss in epithelial tissues and the amount of E-cadherin (Fig. 4F, R^2^=0.875). This suggests that the low amount of cadherin-based junctions in MDA-MB-231 tissues results in a larger resistance to volume loss.

**Figure 4.**
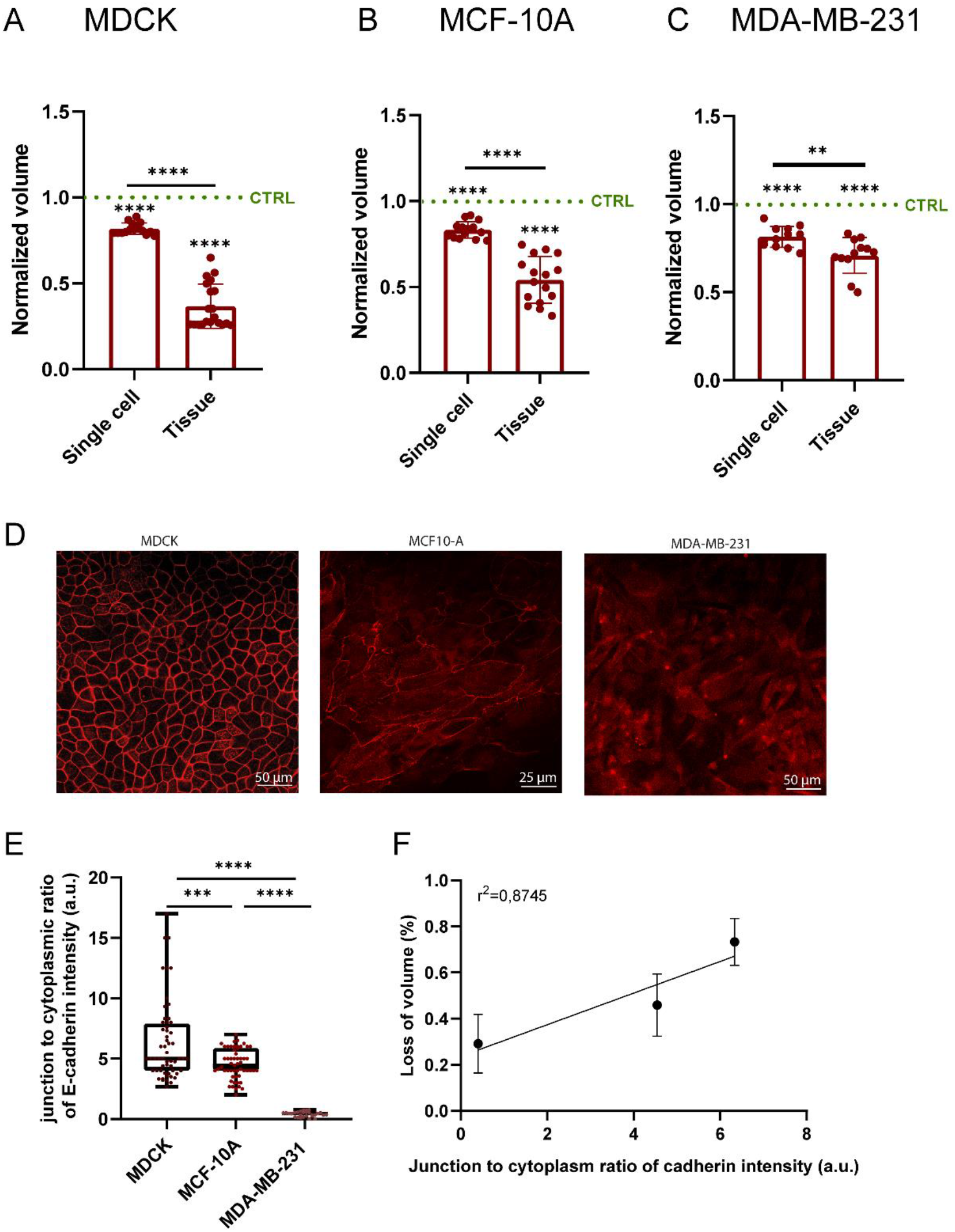
The establishment of cell-cell interactions modulates the volume loss in response to osmotic stress. Comparison of the cell volume of single cells and micropatterned tissues for (A) MDCK, (B) MCF10A and (C) MDA-MB-231 treated with PEG 2.5%. For each cell type, the volumes of PEG-treated tissues are normalized by the volume of micropatterned tissues in isotonic condition. (D) Staining of E-cadherin for MDCK, MCF10A and MDA-MB-231 cells. (E) Quantification of the junction to cytoplasm ratio of E-cadherin intensity for MDCK, MCF10-A and MDA-MB-231 cells. (F) Percentage of volume loss in function of the junction to cytoplasm ratio of E-cadherin intensity. Cells exhibiting higher fluorescence intensity due to cadherin show higher volume loss. n=15 (MDCK single cells), n=20 (MDCK tissues), n=15 (MCF10A single cells), n=16 (MCF10A tissues), n=12 (MDA-MB-231 cells), n=12 (MDA-MB-231 tissues). *0.01<p<0.05, **0.001<p<0.01, ***0.0001<p<0.001, ****p<0.0001 and ns non-significant.

### The effect of osmotic stresses on the migration velocity of epithelial cells is lowered by disrupting cell-cell adhesions

To confirm the role of the cell-cell adhesive interactions on the sensitivity of epithelial tissues to moderate osmotic stress, we perform time-lapse experiments on EGTA-treated tissues. We acquire 8 hours of images under isotonic conditions in culture medium to obtain a control, and then add EGTA solution at 5 mM and acquire an additional 8 hours of time-lapse images. The migration speed increases for MDCK and MCF-10A cells (Fig. 5A-B) due to the disruption of cell-cell adhesions by the EGTA treatment, whereas the migration speed of MDA-MB-231 cells that contain lower amounts of cadherin at cell-cell junctions (Fig. 4E) is not affected by EGTA treatment (Fig. 5C). The effect of EGTA treatment on cadherin disruption is confirmed by immunostaining experiments (Fig. 5D). To further demonstrate the role of cadherin-based adhesions in response to moderate osmotic stresses, we add PEG at 2.5% (v/v) to the EGTA-treated tissues. The PEG treatment does not affect the migration speed of all three cell types (Fig. 5A-C), suggesting that the cadherin-based junctions are more important in controlling the migratory behavior of epithelial cell assemblies under moderate osmotic stress.

**Figure 5.**
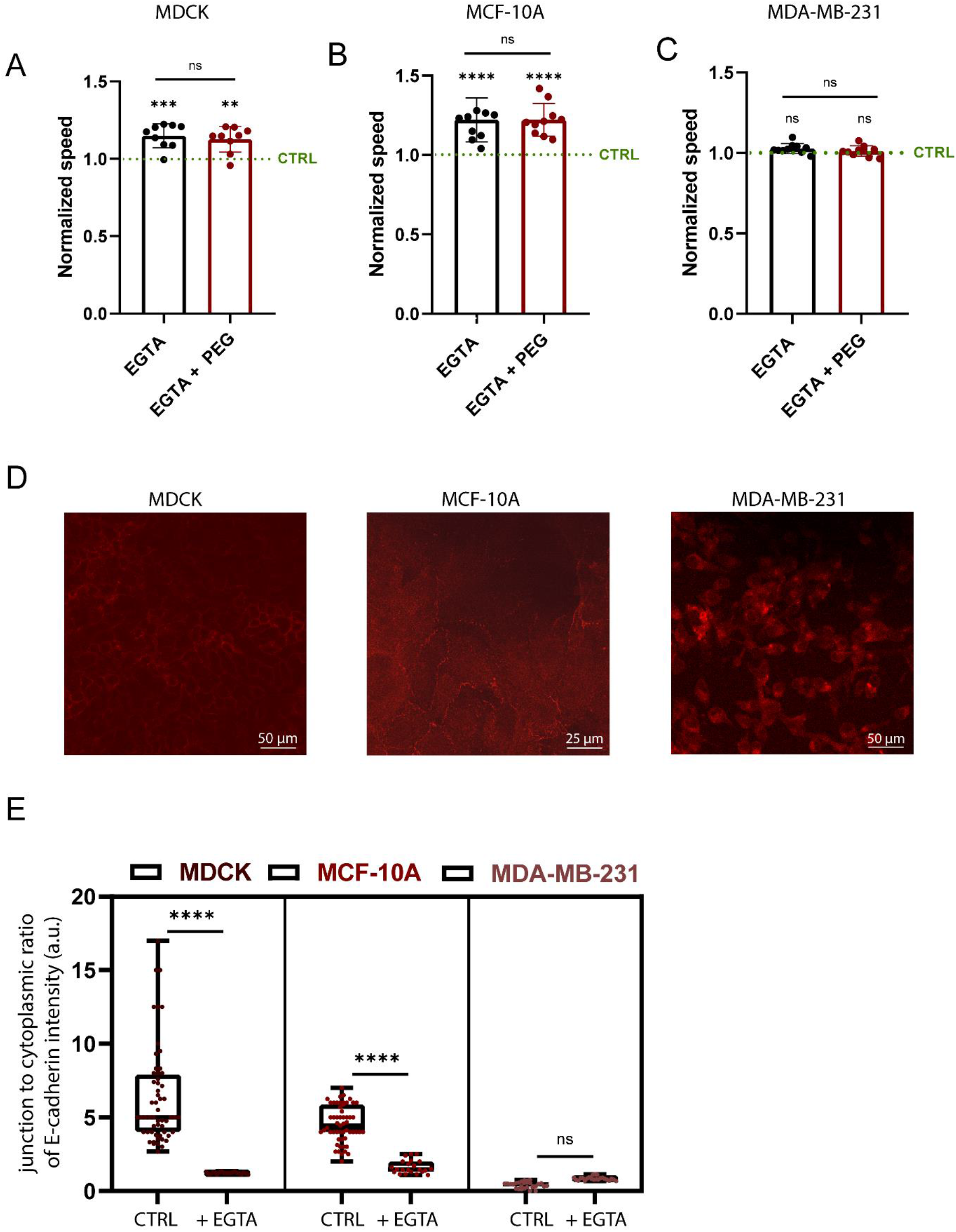
The effect of osmotic stress on the migratory behavior of epithelial cells is reduced by disrupting cell-cell adhesions. (A, B, C) Average speed with EGTA treatment and EGTA treatment following by PEG 2.5% osmotic compression, both normalized by control values (without PEG and EGTA) for MDCK, MCF10A and MDA-MB-231 cell clusters respectively. n=9 (MDCK), n=13 (MCF10A) and n=10 (MDA-MB-231). (D) Immunostaining of E-cadherin after EGTA treatment. (E) Quantification of the junction to cytoplasm ratio of E-cadherin intensity before and after EGTA treatment for MDCK, MCF10-A, MDA-MB-231. *0.01<p<0.05, **0.001<p<0.01, ***0.0001<p<0.001, ****p<0.0001 and n.s. non-significant

## Discussion

Many studies have shown that osmotic pressure can impact cell volume and affect cell structure. (Guo et al., 2017; Miermont et al., 2013; Chowdhury et al., 1992) Osmotic stress can lead to a significant reduction of the cell volume due to water efflux; this results in a corresponding increase in molecular intracellular crowding (Miermont et al., 2013; Mitchison, 2019). However, most of the previous work focused on isolated cells and used high levels of osmotic stress, both of which are far from physiological conditions.

Here, we investigate whether moderate osmotic stresses can affect the migration of breast epithelial cells by using three different epithelial cell lines: a control cell line (MDCK), a non-tumorigenic epithelial breast line (MCF-10A) and a tumorigenic epithelial breast line (MDA-MB-231). By performing OMTC experiments, we find that MDCK cells are stiffer after application of osmotic stress, whereas the mechanical properties of non-tumorigenic and tumorigenic epithelial cells remain constant. In addition, tumorigenic cells are observed to be less sensitive to moderate osmotic stresses than normal and non-tumorigenic cells. In addition, cell volume loss is larger for micropatterned epithelial monolayers than for single cells, highlighting the role of cell-cell junctions.

To demonstrate the role of cell-cell adhesions, we compare the effect of osmotic pressure on cell volumes between single cells and cell clusters. Single epithelial cells are less affected than epithelial cell clusters, suggesting that cell-cell interactions have a key influence on modulation of cell volume in response to osmotic stress. To validate this hypothesis, we perform experiments with EGTA to inhibit cell-cell adhesion in epithelial cell clusters. We find that the cell volume is less affected by osmotic stress in EGTA-treated clusters, regardless of the cell type. This suggests that cell-cell adhesions have a major role in the cell-volume regulation mechanism and hence the sensitivity of epithelial cells to osmotic stress.

To conclude, this study helps unravel the variability in osmoregulatory mechanisms among different epithelial cell lines and highlights the central role of cell-cell interactions. We envision that a better understanding of the osmoregulation of epithelial cancer cells in different *in vitro* conditions might help to better target potential pharmacological agents.

## Material and methods

### Stamp fabrication

Stamps for micropatterning were fabricated by soft lithography using polydimethylsiloxane (PDMS; Sylgard 184 Silicone Elastomer Kit; Dow Corning). Briefly, a thin layer of SU8-3010 (MicroChem) was spin coated onto the surface of a silicon wafer. After baking, a photomask was placed on top of the wafer for UV exposure. Propylene glycol methyl ether acetate (PGMEA) (Sigma-Aldrich, 537543) was then used to remove the unexposed SU8 from the wafer. After baking, the wafer was placed in a petri-dish and served as a mold for downstream PDMS fabrication. PDMS was well mixed with curing agent at a 10:1 ratio, degassed, poured onto the wafer previously passivated for 30 min with fluorosilane (tridecafluoro-1,1,2,2-tetrahydrooctyl-1-trichlorosilane). Finally, the wafer was placed in a 65°C oven for at least 1.5 hours. The PDMS was then cut and carefully peeled off from the mold.

### Cell culture

We used MDCK a renal epithelial cell model, MCF-10A a normal human breast epithelial cell and MDA-MB-231 a human tumor breast cell line. MDCK obtained from ATCC (ATCC CCL-34) were cultured in Eagle’s Minimum Essential Medium with 10% fetal bovine serum and 1% of penicillin/streptomycin at 37°C with 5% CO_2_. MCF-10A cells obtained from ATCC (ATCC CRL-10317) were cultured in MBEM with additives obtained from Lonza/clonetics Corporation as a kit (MEGM, Kit catalog No. CC-3150) and 100 ng/ml of cholera toxin, 1% of penicillin/streptomycin at 37°C with 5% CO_2_. MDA-MB-231 obtained from ATCC (ATCC HTB-26) were cultured in Leitbovitz’s L-15 Medium with 10% fetal bovine serum and 1% of penicillin/streptomycin at 37°C without CO_2_. Cells were allowed to spread on substrates for several hours (>4 h), then were synchronized by starvation in serum-free medium overnight before volume imaging and stiffness measurement. Cells were synchronized to avoid cell size growth and volume changes due to natural variations during the cell cycle (Guo et al., 2017).

### Cell Labeling and Immunofluorescence

Live cells were fluorescently labeled with CellTracker (Invitrogen) SiR-DNA (Sprirochrome) to label cytoplasm and nucleus, respectively. Cells were imaged and sectioned at 0.15-μm intervals by using excitation from a 633- or 543-nm laser or a 488-nm line of an argon laser and a ×63, 1.2-NA water immersion objective or ×40 dry air or ×20 dry air on a laser scanning confocal microscope (TCS SP5; Leica). To label cell-cell adhesions (E-cadherin, Beta-catenin and N-cadherin), cells were previously fixed with paraformaldehyde 4% and permeabilized with Triton-X100 (0.5%) in PBS and stained with 1/200 anti-E-cadherin-FITC (Invitrogen).

### Cell volume

Stained cells were observed by using a ×63/1.2-NA water immersion lens and tissue by using a ×20 dry lens on a Leica TSC SP5. Optical cross-sections were recorded at 0.15-μm z-axis intervals to show intracellular, nuclear, and cortical fluorescence. By using theoretical point spread function, a stack of gray-level images (8 bits) were subjected to deconvolution before 3D visualization. The 3D visualization was carried out by using ImageJ software. The volume was calculated by counting voxel number, after thresholding the stack. Confocal measurements were previously compared with AFM data and the results from the two techniques agreed (Guo et al., 2017).

### Osmotic Stress

Hyperosmotic stresses were applied by adding PEG 300 to isotonic culture medium. The actual osmotic pressure applied to cells was calculated by adding the osmotic pressure due to PEG to that of isotonic solution (330 mOsm/kg) and was further confirmed through a selective measurement by using a micro-osmometer (model 3300; Advanced Instruments, Inc.). Before doing experiment, cells were allowed to equilibrate in PEG solution for 10 min at 37°C and 5% CO_2_. The cell size decreases within 30s and maintains the small volume for hours.

### Confocal microscopy imaging

To observe cell-cell adhesions, cells were seeded onto cell culture membrane assemblies and cultured in a cell incubator (37°C and 5% CO_2_) for 16 hours. Cells were fixed with 4% formaldehyde and 0.1% Triton ×100 diluted in PBS, followed by PBS washes for 3 times to remove excessive reagents. After cell fixation, cells were double stained for actin and E-cadherin or N-cadherin or β-catenin antibody. Fixed cells were blocked with 10% bovine serum albumin (Thermo Fisher Scientific Inc, MA, USA) in PBS for 1 h, followed by a two-step immunostaining process for cadherin. Briefly, cells were first incubated with mouse monoclonal anti-cadherin antibody (Sigma-Aldrich, MO, USA) diluted ×200 in PBS with a supplement of 10% BSA for 45 min at 37°C. Samples were then washed 5 times for 5 min each with PBS and incubated with goat anti-mouse Alexa Fluor 488 secondary antibodies diluted ×200 in PBS with a supplement of 10% normal goat serum for 1 h in the dark. To stain actin, Phalloidin-iFluor 555 (Abcam, MA, USA) are diluted at 1:400 to incubate cells for 1 h. Stained cells are then washed 3 times for 5 min with PBS and imaged with the confocal microscope. Cell-cell adhesions are stained with E-cadherin, N-cadherin and beta-catenin antibody. The image is obtained using ×25 or ×40 water immersion objective on a TCS-SP5 confocal laser scanning microscope (Leica Microsystems Inc., IL, USA).

### Cell tracking

Time-lapse microscopy experiments were performed with the Matlab pluggin Celltracker ^46^ and analyzed with Prism Graphpad. The cell nucleus was stained with SirDNA (Spirochrome) to allow time-lapse microscopy experiments in live conditions for at least 16 hours. We used a Leica TSC SP5 microscope equipped with ×20 magnification lens.

### Stiffness Measurements

The cell mechanical properties were probed by using OMTC, which is a high-throughput tool for measuring adherent cell mechanics with high temporal and spatial resolution. Measurements were performed at 37°C. In brief, functionalized ferrimagnetic beads (4.5 μm) coated with PLL (4 kDa) were incubated with cells for 20 min in the incubator. For stiffness measurements, beads were first magnetized horizontally by a large magnetic field. A much weaker magnetic field (oscillating at 0.75 Hz) was then applied vertically, which applied a sinusoidal torque to the beads. Motion of the beads was optically recorded. The ratio between the torque and bead motion thus defined an apparent cell stiffness (Pa/nm). A series of geometric factors, based on finite element models that take into account the cell thickness and bead-cell coupling, can be used to convert the apparent stiffness into shear modulus of the cell.

### E-cadherin Contrast Calculation

On the E-cadherin immunofluorescence images, an average intensity was measured at the cell−cell junction (i_jct_) and in the cytoplasm (i_cyt_) using ImageJ. These values where averaged over 20-25 cells (I_jct_=〈i_jct_〉 and I_cyt_=〈i_cyt_〉). We then defined J, the junction to cytoplasm ratio of E-cadherin intensity : J=I_jct_/I_cyt_.

## Acknowledgment

D.M. acknowledge the financial support of the Belgian American Education Federation (BAEF) and the Fédération Wallonie Bruxelles (WBI). The authors thanks John Heyman, Jing Xia and Sijie Sun for their advice. The work at Harvard was supported by the NIH (P01HL120839; 1R01HL148152, 1U01CA202123). The authors declare no competing financial interests.

## Authors contributions

D.M. and D.A.W. conceived the project and D.A.W. supervised the project. D.M. and P.Y.C. performed the experiments. D.M. analyzed the data and wrote the main manuscript text and prepared figures. D.M., D.A.W., P.Y.C. and J.J.F. contributed to the interpretation of the results and improved the manuscript and figure presentations.

---

**Supplementary 1.**
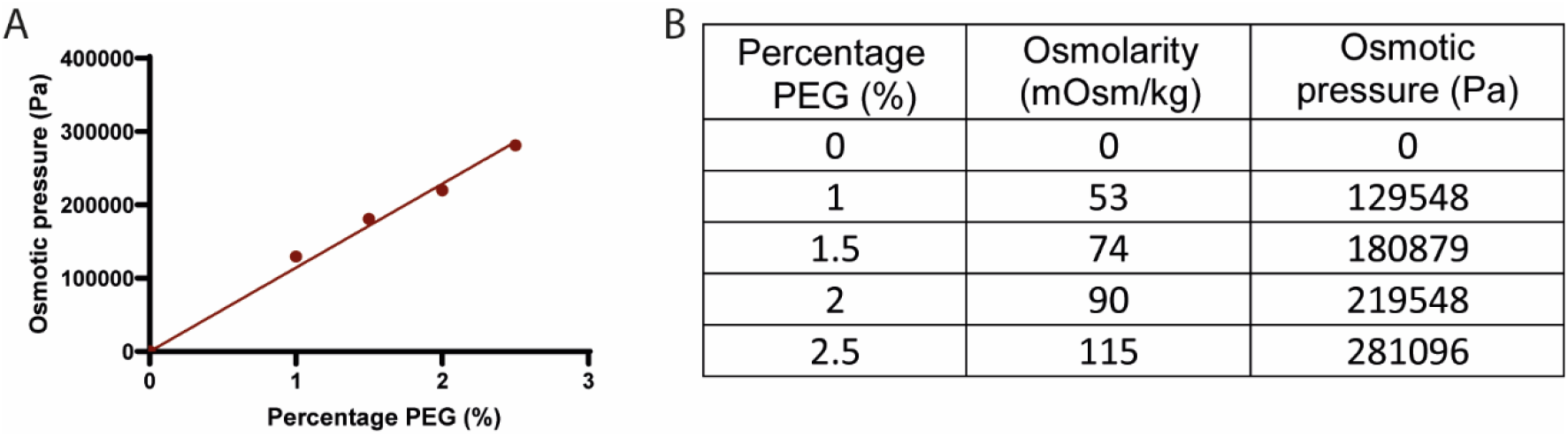
(A) Calibration curve of osmotic pressure measured in function of PEG concentration. (B) Table with values of osmolarity measured with an osmometer and the corresponding osmotic pressure.

